# Modulation of Human Hsp90α Conformational Dynamics by Allosteric Ligand Interaction at the C-Terminal Domain

**DOI:** 10.1101/386755

**Authors:** David L. Penkler, Özlem Tastan Bishop

## Abstract

Recent years have seen heat shock protein 90 kDa (Hsp90) attract significant interest as a viable drug target, particularly for cancer. To date, designed inhibitors that target the ATPase domain demonstrate potent anti-proliferative effects, but have failed clinical trials due to high levels of associated toxicity. To circumvent this, the focus has shifted away from the ATPase domain. One option involves modulation of the protein through allosteric activation/inhibition. Here, we propose a novel approach: we use previously obtained information via residue perturbation scanning coupled with dynamic residue network analysis to identify allosteric drug targeting sites for inhibitor docking. We probe the open conformation of human Hsp90α for druggable sites that overlap with these allosteric control elements, and identify three putative natural compound allosteric modulators: Cephalostatin 17, 20(29)-Lupene-3β-isoferulate and 3′-Bromorubrolide F. We assess the allosteric potential of these ligands by examining their effect on the conformational dynamics of the protein. We find evidence for the selective allosteric activation and inhibition of Hsp90’s conformational transition toward the closed state in response to ligand binding and shed valuable insight to further the understanding of allosteric drug design and Hsp90’s complex allosteric mechanism of action.

## Introduction

The 90 KDa heat shock protein (Hsp90) is a highly conserved molecular chaperone crucially involved in maintaining cellular homoeostasis in organisms from most kingdoms of life with the exception of archea^1^. In the cytosol, Hsp90’s main biological function is the facilitation of folding, maturation, and trafficking of numerous client peptides both native and denatured^2–4^. Hsp90’s diverse array of clientele implicate the chaperone in several associated biological functions and place it at the intersection of various fundamental cellular pathways, where it acts as a central hub in maintaining numerous protein interaction networks^1^.

Hsp90 exists as a homodimer (**Figure 1-A**), and each protomer is comprised of three well characterized domains^5–7^: an N-terminal domain (NTD) which is responsible for ATPase activity and facilitating transient inter-protomer dimerization^8^; a middle domain (M-domain) that provides a large surface area for cofactor and client binding and contributes to ATPase activation^9^; a C-terminal domain (CTD) which serves as the primary site for inter-protomer dimerization^10,11^. The NTD and M-domain are connected by a highly flexible charged linker that has been implicated in modulating chaperone function^12–15^. Hsp90’s molecular function critically hinges around its ability to bind and release client peptides via a complex nucleotide dependent conformational cycle (**Figure 1-B**). In a nucleotide free state, the dimer becomes highly flexible and is capable of assuming multiple conformers with a higher affinity for an open “v-like” conformation in which the M-domains of each protomer are suitably exposed for client loading^16–18^. ATP binding triggers structural rearrangements in the NTD that promote dimerization at the N-terminal, stabilizing a closed catalytically active conformation^10,19^. Transition to the closed ATPase active state is an inherently slow process recording time constants in the order of minutes^8,20,21^, possibly due to energetic barriers presented by structural intermediates that may be overcome through cofactor mediation^22–25^. ATP hydrolysis and the subsequent release of ADP from the NTD initiate a conformational return to the native apo open state and client release.

**Figure 1:**
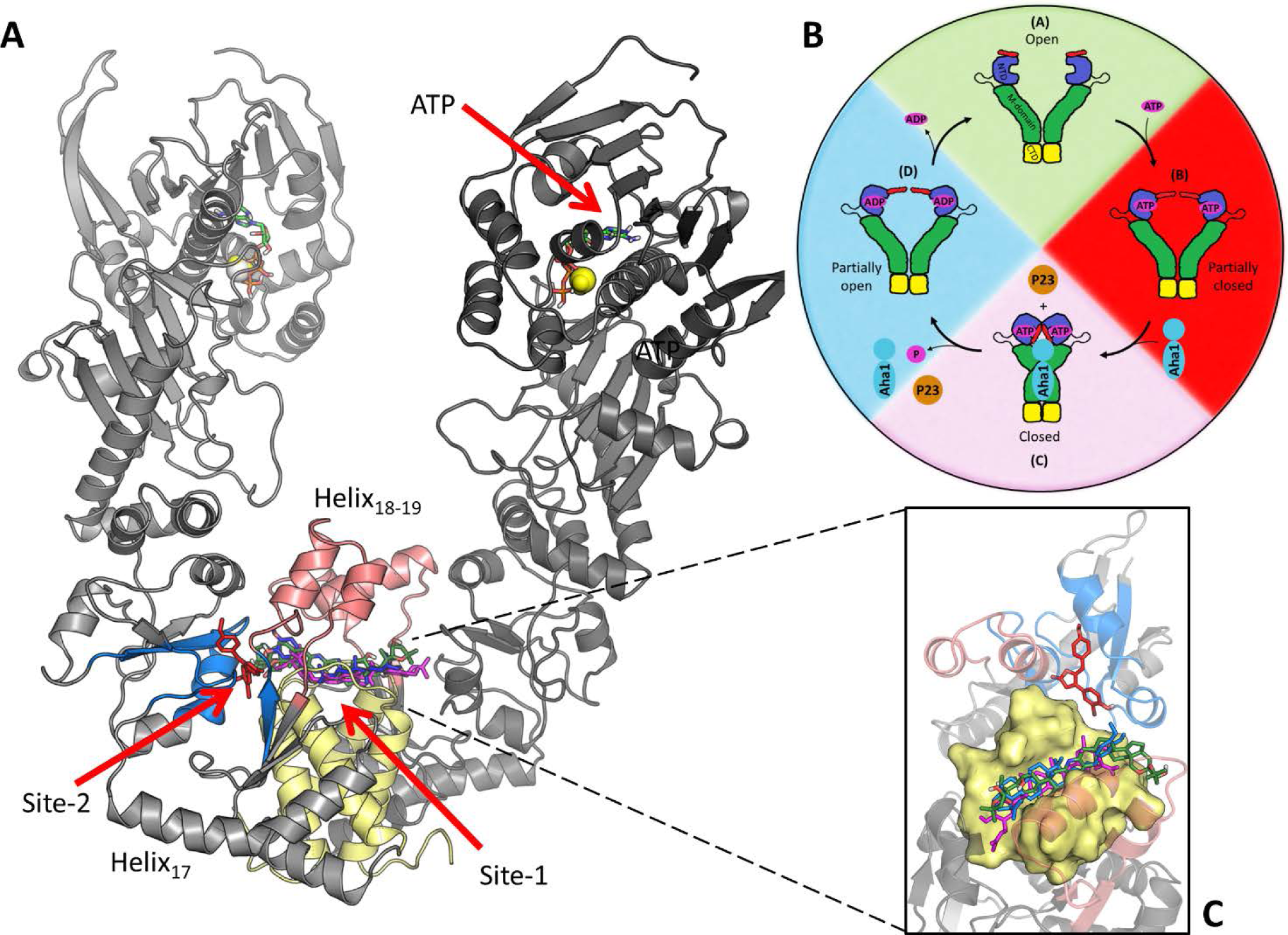
Illustration of Hsp90α in the open conformation. **(A)** The location of the different binding site residues are shaded: Site-1 helix_18-19_ (red), helix_21-22_ four-helix bundle (yellow) and Site-2 sub-pocket (blue). The NTD location of ATP and magnesium ions (spheres) are also shown. (**B**) Hsp90’s nucleotide driven conformational cycle (Adopted from Penkler et al.^37^). (**C**) Inset – zoomed in view of docked compounds SANC309 red, SANC491 green, SANC518 blue, and Novobiocin magenta.

The interaction of Hsp90 with several disease related peptides implicate the chaperone with the progression and development of several associated pathologies such as protein folding disorders, cancer, and neurological disease^5^. In recent years it has become increasingly clear that deregulation of Hsp90 may present an attractive treatment strategy for these diseases, elevating interest in human Hsp90 as a viable drug target particularly for the treatment of cancer^26–28^. To date, Hsp90 inhibitors have largely targeted the ATPase domain, demonstrating potent anti-proliferative effects; however, their progression through clinical trials has been severely limited due to high levels of toxicity associated with an induced heat shock response^26^. For this reason, recent research has shifted focus away from the NTD to targeting distant functional sites of the protein as alternative drug treatment strategies. The design and development of inhibitors that target the formation of specific Hsp90-co-chaperone complexes, has led to the discovery of inhibitors such as Derrubone, Withaferin A, and Celastrol which block the Hsp90-Cdc37 complex^29^. Discovery of a secondary nucleotide binding site at the CTD led to the design and development of several CTD inhibitors including the coumarin based antibiotic Novobiocin that has been shown to allosterically destabilise the Hsp90 dimer by interfering with CTD dimerization leading to the dissociation of bound clients^30–32^. These CTD inhibitors of Hsp90 have a distinct advantage over the NTD ATPase inhibitors, in that they do not appear to induce secondary heat shock responses and thus incur lower levels of toxicity^29^.

To date, biochemical and computational studies have demonstrated allosteric coupling between the NTD and CTD^33–37^ suggesting nucleotide driven conformational restructuring may be influenced by allosteric events at the CTD. More specifically, computational studies based on the closed conformation of yeast Hsp90 have identified a putative allosteric binding site located at the M-domain:CTD inter-protomer interface that is implicated in allosteric modulation of conformational dynamics^33,35,38^. Subsequent to this discovery, further studies have led to the development of several allosteric CTD ligands that appear to enhance ATPase activity up to six-fold by promoting conformational dynamics in favour of the ATPase active closed conformation^39,40^. Furthermore, a recent biochemical study has provided evidence of Bisphenol A based CTD inhibitors of human Hsp90α that bind a site separate from the nucleotide binding site^41^ and display anti-proliferative activity in tumour cell lines^42^. In a previous computational study, we used all-atom molecular dynamics (MD) simulations coupled with dynamic residue networks (DRN) and perturbation response scanning (PRS) to determine allosteric hotspots that may be implicated in modulating conformational dynamics in human Hsp90α^37^. Focusing on the open “v-like” conformation, we identified several residues located at the M-domain:CTD interface that form a central hub in the protein interaction network, and provided evidence that when externally perturbed by random force displacements these residues are capable of selecting global conformational displacements towards the ATPase active closed conformation^37^.

In the present study we propose that force perturbations at the aforementioned sites, may occur naturally through binding forces engendered by protein-protein and protein-ligand interactions; and that these allosteric hotspots may present a suitable allosteric drug target candidates. We probe two putative ligand binding sites that were identified by combined PRS and DRN analysis^37^, and supported by FTMap^43^ screening at the CTD as potential allosteric drug targeting sites (**Figure 1** Site-1 & Site-2). We found three South African Natural compounds (SANC) that dock to these sites: Cephalostatin 17 (SANC491) and 20(29)-Lupene-3β-isoferulate (SANC518) preferentially bound at Site-1 together with the known CTD inhibitor Novobiocin which was included for comparative control purposes; and 3′-Bromorubrolide F (SANC309) docked at Site-2. We utilized several all-atom MD simulation based analysis techniques, to assess the allosteric potential of each site to modulate protein conformational dynamics in response to ligand interactions.

## Results and Discussion

### Identification of novel natural compounds that bind the CTD

The availability of putative small molecule binding sites at the CTD of the “v-like” open conformation of human Hsp90α^37^ was assessed using FTMap^43^ with an unbiased screen positioned over the M-domain:CTD region, resulting in two candidate binding sites overlapping with allosteric control elements and identified in our previous study via combined PRS and DRN analysis^37^ (**Figure S1**). Site-1 is at the CTD interface in a groove positioned at the four-helix bundle (helix_21,22_) (**Figure 1**-yellow). Site-2 is adjacent to Site-1 and represents a sub-pocket positioned at the M-domain:CTD interface of each protomer (**Figure S1 –** 2A & 2B). Each sub-pocket is formed by residues belonging to helix18, ß-sheet_17-18_, and helix_22_, as well as several residues located at the M-domain:CTD hinge region.

Molecular docking was utilized to separately screen a library of 702 South African natural compounds against Site-1 and Site-2. The known CTD inhibitor Novobiocin was included in both screens as a control. Due to computational constraints, the protein receptor was kept rigid and the ligands allowed flexibility around available rotatable bonds. Each compound was re-docked 100 times to ensure conformational diversity and the results filtered and analysed in terms of conformation cluster size and average estimated free energy of binding. Table 1 provides a summary of the best docked compounds for each site.

**Table 1:** Summary of the best docked compounds for Site-1 and Site-2

| Compound | Name | Site | Cluster size | Average energy (kcal/mol) | Lowest energy (kcal/mol) |
| --- | --- | --- | --- | --- | --- |
| Novobiocin | Novobiocin | 1 | 81 | -8.75 | -8.97 |
| SANC309 | 3'-Bromorubrolide F | 2 | 79 | -9.13 | -9.20 |
| SANC491 | Cephalostatin 17 | 1 | 93 | -11.41 | -11.44 |
| SANC518 | 20(29)-Lupene-3β-isoferulate | 1 | 84 | -11.03 | -11.46 |

Of the 702 natural compounds, only three putative hits against either CTD binding site were observed (Table 1, **Figure 2**). Novobiocin, Cephalostatin 17 (SANC491) and 20(29)-Lupene-3β-isoferulate (SANC518) reproducibly docked to Site-1. All three ligands recorded similar docking orientations, each compound lining the length of a binding groove positioned over the four-helix bundle with their terminal moieties extending towards the Site-2 sub-pocket in protomer B (**Figure 1**-inset). This observed binding site and orientation is in agreement with previous reports for Novobiocin^44^. Both SANC491 and SANC518 recorded lower binding free energy scores (<-11.0 kcal/mol) compared to Novobiocin (−8.97 kcal/mol) suggesting higher binding affinities. Looking at Site-2, only 3′-Bromorubrolide F (SANC309) was able to dock to the small sub-pocket in protomer B with a high degree of reproducibility and suitable binding free energy scores (−9.13 kal/mol) (**Figure 1** – red). To fully examine the stability of ligand binding and to gain further insights as to the allosteric effect of each ligand, all-atom MD simulations were carried out on the protein-ligand complexes for a total 200 ns. In each case, the ligand conformation with the lowest binding free energy score from the largest cluster was selected as a representative start structure.

**Figure 2:**
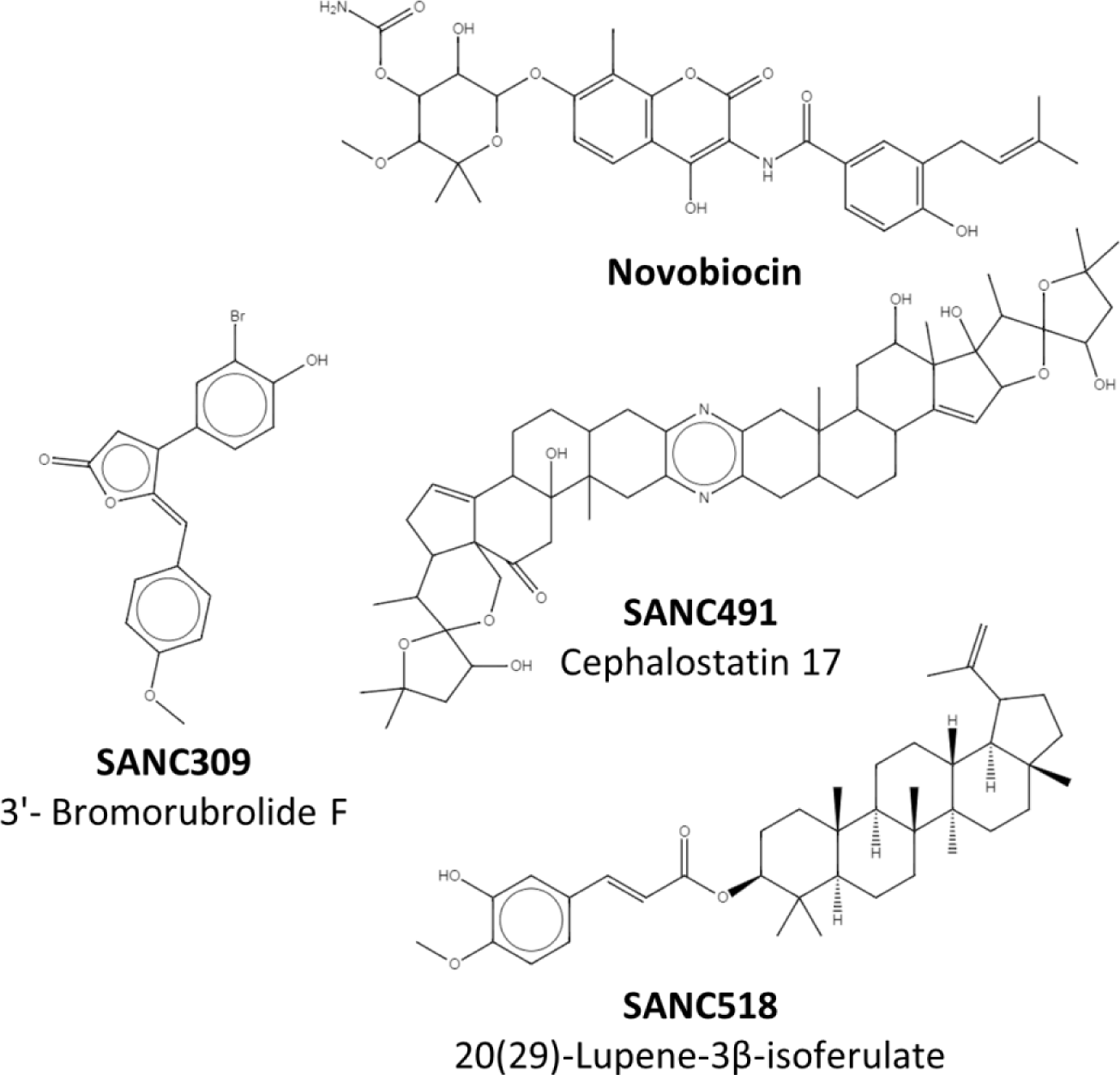
2D representation of Novobiocin and the identified natural compounds that bind the C-terminal.

### Characterization of protein-ligand interactions

Ligand binding stability was assessed by monitoring the residue contribution to protein-ligand interactions over time. Hydrophobic and hydrogen bond interactions were evaluated every n^th^ frame in the trajectory using a 200 ps time interval, and the cumulative data represented as interaction heat maps (**Figure 3**). Starting with binding Site-1, Novobiocin, SANC491, and SANC518 primarily interact with residues (residues L672-Q682) from both protomers that line a distinct binding groove located at four-helix bundle of the CTD inter-protomer dimerization interface (**Figure 3** – yellow). Of these residues, L627, S677, L678, and P681 are the most consistent contributors to ligand interactions. In addition to stable hydrophobic (residues L627 and P681) and hydrogen bond interactions (residues S677 and L678), the terminal moieties of the Site-1 ligands also interact with residues (T495-F507) that line the entrance to the sub-pocket binding site in protomer B (Site-2) (**Figure 3** – blue). The only notable difference between the Site-1 ligands was the formation of additional interactions between SANC491 and helix_18_ of protomer A (residues R612-T624) (**Figure 3** – red). The observed orientation of the binding Site-1 ligands together with their corresponding interacting residues is in close agreement with a previous biochemical and *in silico* study of Bisphenol A based allosteric inhibitors of human Hsp90^42^. Furthermore, interacting residues L672, S674, and P681 are closely positioned to and overlap with CTD allosteric hotspots (residues599-W606, and T669-L678) previously implicated in NTD allosteric signalling and control of conformational dynamics^33^.

**Figure 3:**
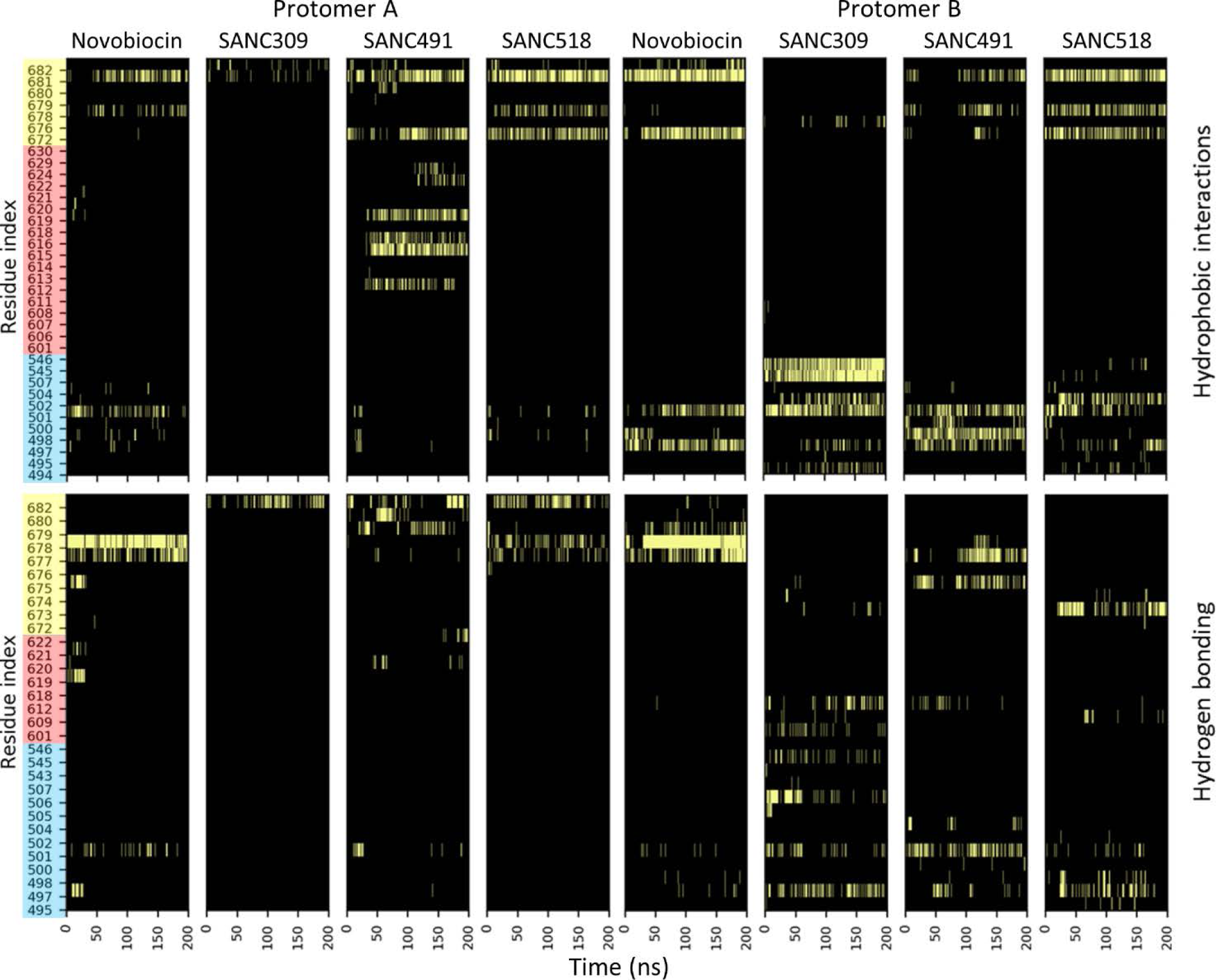
Time evolution of residue contribution to protein-ligand hydrophobic and hydrogen bond interactions. Detected interactions are depicted by light bars. Y-axis residue shading represents the different binding site residues: blue - sub-pocket; red – helix_18_; yellow - four-helix bundle.

Looking at binding Site-2, SANC309 appears to interact exclusively with residues belonging to protomer B (residues T495-F507 and S543-K546, **Figure 3** – blue) with the exception of hydrogen bond interactions with the four-helix bundle through residue Q682 in protomer A (**Figure 3** – red). In protomer B, residues Q501, T545 and K546 form stable hydrophobic interactions with SANC309 while interactions with the remaining sub-pocket residues appear to be more transient (**Figure 3** – blue). The protein-ligand interaction landscape observed for SANC309 is to the best of our knowledge novel to the current study and notably overlaps with several allosteric hotspot residues (T495, E497, T545, and K546) that have been previously implicated in selecting conformational displacements in favour of the closed conformation when externally perturbed^37^.

Overall, MD simulations revealed stable protein-ligand complexes over 200 ns, and the interaction profiles for both Site-1 and Site-2 overlap with known allosteric sites opening the possibility for external modulation of Hsp90α conformational dynamics through ligand binding interactions. We investigate this possibility by monitoring the effect each ligand has on the global and internal dynamics of the protein compared to a ligand-free system and assess the respective allosteric potential of Site-1 and Site-2.

### CTD ligands modulate conformational sampling and protein flexibility

Backbone root-mean-square-deviation (RMSD) analysis serves as a measure for monitoring conformational change over an MD trajectory. In the absence of bound ligand, whole protein RMSD analysis indicates variable backbone flexibility with RMSD values around 0.65 nm (**Figure S2**-black). This observation is not surprising given the inherent plasticity of Hsp90 in its open conformation^45^. Addition of Novobiocin or SANC518 appears to improve the stability of the dimer compared to the ligand-free system, recording RMSD values around 0.50 nm. In contrast, the SANC309 and SANC491 complexes appear to experience more variable conformational sampling, with RMSD values ranging between 0.35 nm and 1.35 nm. This data tentatively suggests an increased flexibility for the latter two compounds and a maintained if not reduced flexibility for Novobiocin and SANC518. The RMSD profiles for each protomer in isolation (**Figure S2**-red, blue) reveal stable protomer backbone conformations for each complex with values around 0.25nm. This excludes major conformational restructuring within the individual protomers as the main cause for the observed dimer flexibility. It is likely that the observed backbone flexibility/mobility may result from conformational repositioning of the rigid protomers relative to one another in either a linear opening/closing manner, or through perpendicular rotation about the C-terminal dimerization axis.

The distribution of the inter-protomer distance, defined as the measured distance between the center of mass for each NTD, informs on the propensity for Hsp90 to populate conformations distinct from the starting structure over the course of the MD trajectory (**Figure 4**). In the absence of CTD ligands, the distribution is positioned around the initial NTD-NTD distance of 7.6 nm (**Figure 4** – black), indicating limited conformational variation. For the ligand bound systems, Novobiocin and SANC518 appear to populate conformations analogous to the ligand-free system (**Figure 4** – magenta & blue), while SANC309 and SANC491 cause a shift in the distribution away from the ligand-free system. SANC491 populates conformations with an increased inter-protomer distance of ~8.5 nm (**Figure 4** – green), suggesting a conformational preference for a more open “v-like” structure. SANC309 on the other hand, populates conformations in favour of the closed conformation recording a distribution peak around 6.7 nm.

**Figure 4:**
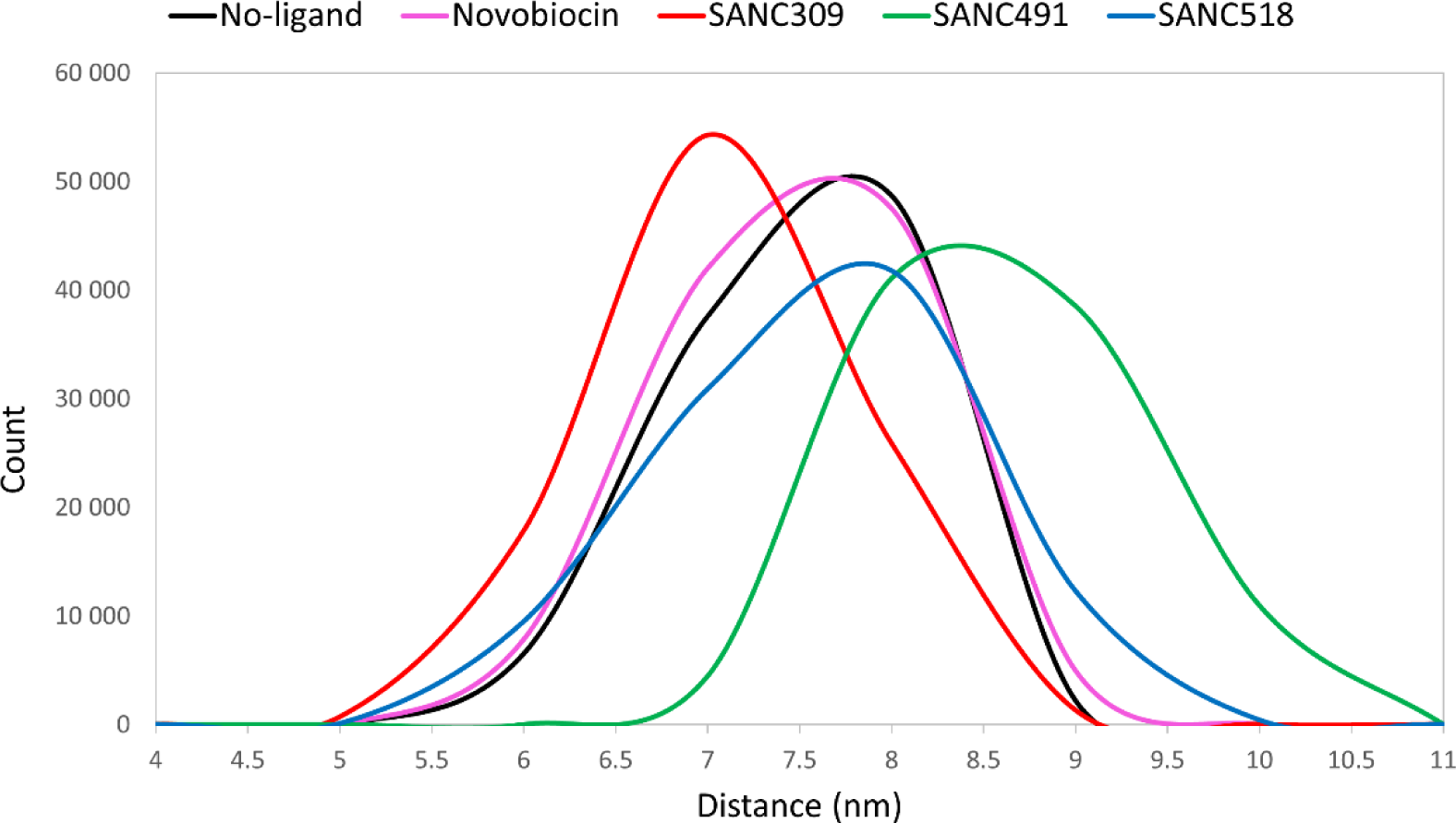
Distribution of inter-protomer distance. Inter-protomer distance is defined as the distance between the center of mass of each NTD.

The distribution of the NTD-CTD distance provides an indication of protomer flexibility around the central axis as seen in hinge bending motions. Shorter NTD-CTD distances could be indicative of protomer bending, while increased NTD-CTD distances could suggest protomer straightening/extension. The NTD-CTD distance is defined as the distance between the centers of mass of the NTD and CTD four-helix bundle (**Figure 1**-yellow), and the time evolution of these measurements is represented as a distribution for each protomer separately (**Figure 5**). In the absence of bound ligand, both protomers form distinct distribution peaks that correspond to the initial measurements of 9.0 nm and 8.0 nm suggesting minimal bending motion and confirming the rigid nature of the protomers under these conditions. Addition of Novobiocin appears to have a minimal effect on conformational sampling, with analogous distributions to the ligand-free complex. Introduction of the natural compounds, on the other hand, appears to skew the distribution curves in both protomers over a wider range of distances, suggesting increased flexibility around the central axis particularly for protomer B. All three compounds shift the distribution peaks in protomer B by 0.5-1.0 nm, while more defined distributions around 9.0 nm are observed for protomer A. The increased NTD-CTD distances in this protomer may suggest favoured sampling of a more extended protomer.

**Figure 5:**
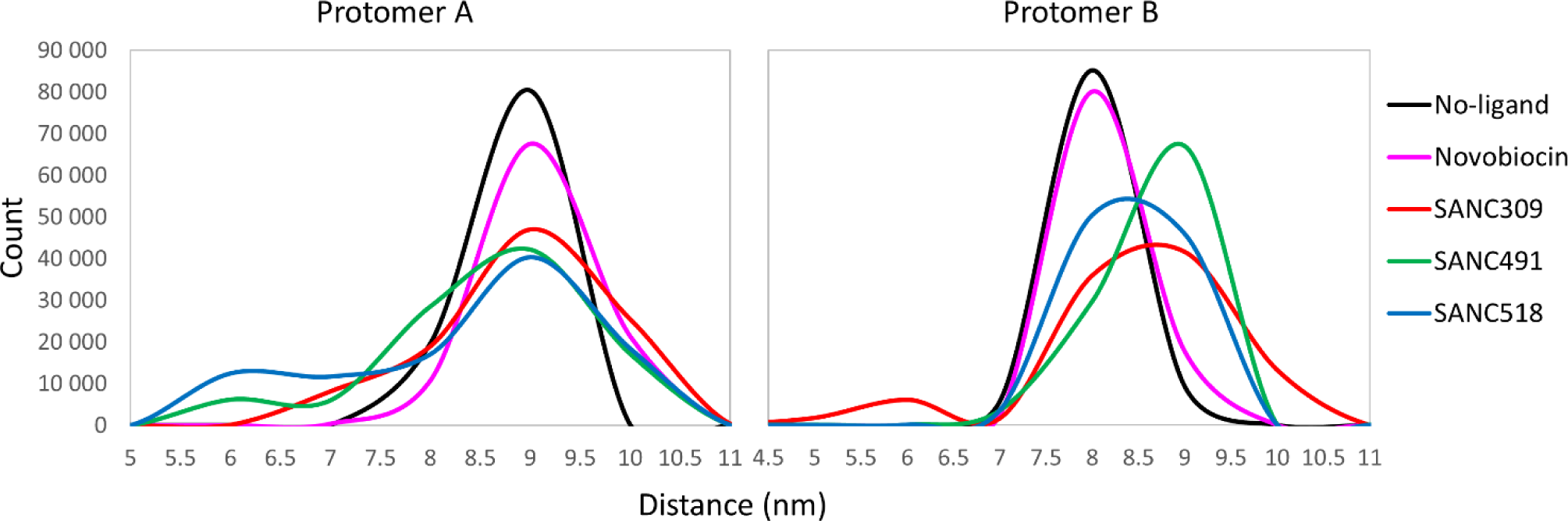
Flexing around the central axis of each protomer. Protomer flexing is measured as the distance between the center of mass of each NTD and the CTD interface.

Next, we focus on the relative effect of bound ligand on the flexibility of localized regions of the protein. The root-mean-square-fluctuation (RMSF) is a measure of the average positional displacement of each residue over time and indicates flexibility/mobility of residue sites. Here, we monitor the relative change in residue fluctuation (ΔRMSF) between the ligand-free and ligand bound complexes to determine the extent to which bound ligands influence intra-protomer flexibility. The ΔRMSF profiles (**Figure 6**A) reveal differential ligand specific modulation of domain flexibility, whereby bound ligands appear to modulate the RMSF of entire domains rather than individual residue sites. Novobiocin and SANC491 differentially modulate the RMSF of the NTDs of each protomer, causing increased fluctuations in protomer A and a decrease in protomer B (**Figure 6**A – blue shading). Meanwhile, SANC309 and SANC518 appear to increase residue fluctuations at the NTDs of both protomers recording large positive ΔRMSF. In addition, SANC309 also experiences increased fluctuations in the M-domains of both protomers compared to the Site-1 ligands (**Figure 6**A – green shading).

**Figure 6:**
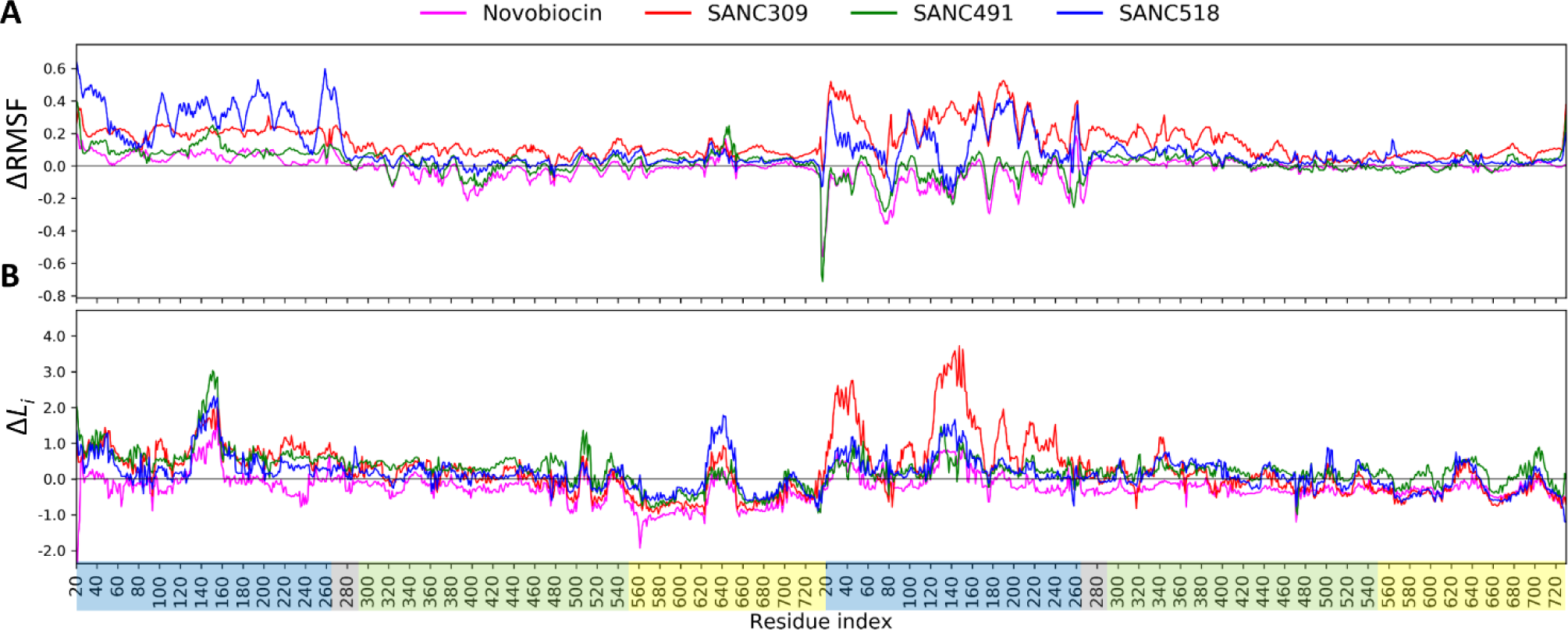
Effect of CTD bound ligands on internal dynamics. (**A**) ΔRMSF, and (**B**) Δ*L*_*i*_ plots for each ligand bound complex, calculated as the average difference between the protein-ligand and ligand-free complexes for each residue. Residue indices are coloured by domain: NTD – blue, M-domain – green, CTD – yellow.

Taken collectively, the results demonstrate evidence of selective ligand modulation of both whole protein mobility and domain specific flexibility depending on the identity of the bound ligand. Conformational flexibility and rigid body mobility form the basis of enzymatic catalysis and allosteric modulation^46^ and in the case of Hsp90, conformational plasticity is crucial for molecular functionality^5,45^. The increased flexibility experienced by SANC309, especially in the M-domain where co-chaperones bind, may enable the protein to overcome energetic limitations allowing it to explore a larger conformational space, thus aiding its search for the closed catalytically active state. Conversely, reduced protomer flexibility in SANC491 may lead to protomer elongation and a more rigid structure that favours the open conformation.

### Ligand modulation of residue interaction networks

Dynamic residue networks (DRNs)^47^ were utilized to analyse the effect of ligand binding on residue connectivity over time. DRNs were constructed for each MD trajectory by treating C_ß_ atoms (C_α_ for glycine) as nodes in the network and connections between nodes established based on a distance cut-off of 6.7 Å (see Methods for details), and the resultant DRNs analysed in terms of average long-range residue reachability (*L*) and average betweenness centrality (BC). In graph theory, the reachability of a residue is defined as the number of connections required to reach residue *i* from *j* using the shortest possible path. The average reachability of a residue (*L*_*i*_) is thus defined as the average number of steps required to reach residue *i* from any other residue in the network. The metric BC is related to *L* in that it is a measure of how often on average a residue is utilized in shortest path navigation and has been previously shown to be an effective measure for the identification of functional residues implicated in intra-protein communication^37,48^, protein-ligand^49^ and protein-protein binding sites^50,51^, as well as nonsynonymous SNP analysis^52,53^. To evaluate the relative effect of CTD bound ligand on the connectivity of the protein we compare each protein-ligand complex to the ligand-free ATP-only complex by monitoring the change in *L*_*i*_ (Δ*L*_*i*_) and BC (ΔBC) over the course of the MD trajectory.

Starting with Δ*L*_*i*_ (**Figure 6B**), it is evident that ligand binding leads to an increase in *L*_*i*_, particularly at the NTDs of both protomers (~1 unit), with the largest increase observed for residues belonging to ATP-lid (residues 130-160). This observation is notably accentuated in protomer B of the SANC309 complex, with Δ*L*_*i*_ values >3.0. It has been previously shown that Δ*L*_*i*_ can be influenced in one of two ways: (i) a significant spatial alteration in the local neighbourhood surrounding *i* can affect the contribution of first and second neighbours to the average number of steps taken^37,54^; (ii) the local network surrounding *i* remains intact but large conformational changes at distant locations in the protein affect the average path length. We have previously shown that there exists a proportional relationship between RMSF and *L*_*i*_, when residue fluctuations/displacements exceed the distance threshold used to construct the DRN^37^. This observation is evident in the present study, RMSF and *L*_*i*_ recording strong Pearson’s correlation coefficients (> 0.85) for all five complexes (**Figure S3**A). Interestingly, this relationship does not hold true when considering ΔRMSF and Δ*L*_*i*_, which record significantly poorer Pearson’s correlations (< 0.7) (**Figure S3**B). This observation suggests that the positive Δ*L*_*i*_ experienced in all the ligand bound systems is not directly linked to increased residue fluctuations as described in scenario (i), and thus we consider scenario (ii) for an explanation for the increase in *L*_*i*_.

The variable flexing/bending of both protomers in response to bound ligand (**Figure 5**), may result in sufficient conformational deformations so as to disrupt established key interdomain network contacts and concomitantly increase the average path length for entire regions of the protein. To investigate this possibility we consider the time progression of all NTD:M-domain contacts using a 200 ps time interval. Inter-domain contacts that maintain a spatial distance ≤ 6.7 Å are colored according to percentage contact duration over time (**Figure 7B**). Looking at the ligand-free system, it is evident that there are more inter-domain contacts between the NTD and M-domain in protomer A compared to protomer B, however two key inter-domain contact regions are present in both protomers (**Figure 7A**): residues K204, V207, I214, I218, and L220 in the NTD form stable contacts with L290 and N291 over 200 ns. In addition to these contacts, residues D57, R60, Y61 and L64 in the NTD of protomer A maintain stable inter-domain interactions with R366-V368. For the ligand bound systems, it is evident that the inter-domain interactions through reside L290 are maintained in both protomers, and those involving N291 are weakened in protomer B. Notably, the stable inter-domain interactions between the R366-V368 triad in protomer A and the NTD are lost in all ligand bound complexes with the exception of SANC491 which retains partial interactions between R367 and the NTD residues D57 and R60. Taken together this suggests that breakdown of stable inter-domain interactions, particularly those involving the R366-V368 triad in protomer A and N291 in protomer B, may lead to NTD:M-domain decoupling, which in turn may result in the redirection of inter-domain network communication to pass through the long NTD:M-domain linker and ultimately increase *L*_*i*_. Indeed, NTD decoupling may be evidenced by the enhanced NTD residue fluctuations observed at the NTD of protomer A (**Figure 6A**).

**Figure 7:**
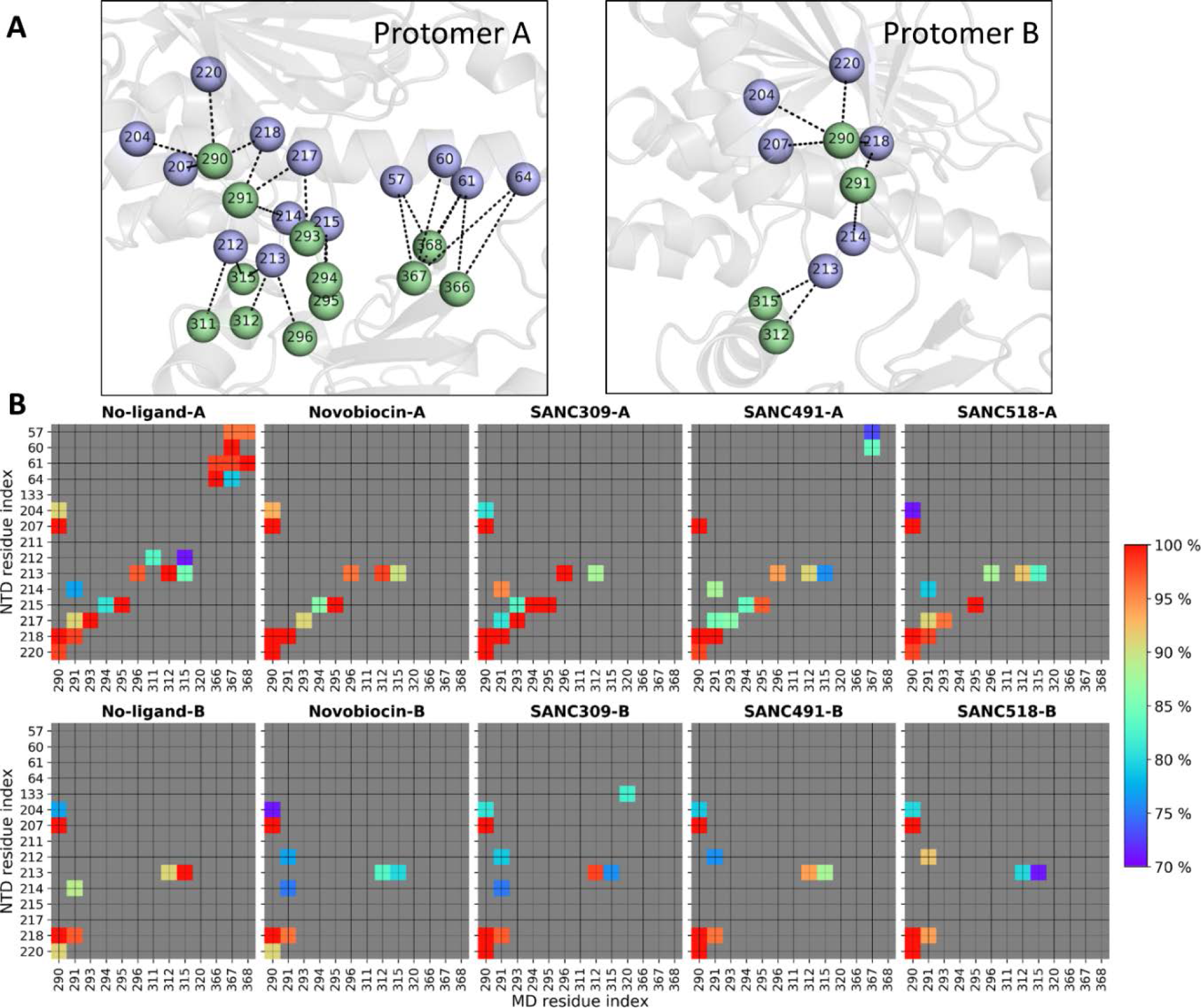
Inter-domain contacts between the NTD and M-domain of each protomer. **(A)** Illustration of the inter-domain contacts between the NTD (violet spheres) and M-domain (green spheres) for the ligand-free ATP-only complex. (**B**) Time progression of key contacts between the NTD and M-domain over the respective MD trajectory for each complex. Contacts are colored (blue – red) by percent detected over the trajectory and grey if detected < 70 %.

While the NTD and M-domain record elevated Δ*L*_*i*_, the CTDs experience a ~1 unit decrease in *L*_*i*_ in both protomers. This observation is particularly apparent for residues 550-620 belonging to helix_17_ and helix_18_, as well as the terminal four-helix bundle (residues 660-720). Interestingly, the former have been previously implicated as sensors to physical perturbations elsewhere in the protein^37^ and implicated in signal propagation^33^. Here, scenario (i) provides a suitable explanation in that ligand binding at the CTD stabilizes the domain reducing residue fluctuations thus allowing for stable residue-residue interactions.

As with *L*_*i*_, ΔBC is also averaged over time and comparison of the different ligand bound complexes reveals a high degree of overlap between the ligand binding sites and regions of the protein that experience large shifts in BC (**Figure S4**), demonstrating the metric’s sensitivity to putative ligand binding sites. Interestingly, all four ligand complexes experience increased BC at the four-helix bundle in protomer A, but a decrease in protomer B. Furthermore, it is evident that the ligand interactions between SANC491 and helix_18_ (red) in protomer A has a direct impact on BC at this location possibly due to ligand stabilisation of this inherently flexible region as demonstrated by the differential Δ*L*_*i*_ values (**Figure 6**). Indeed, this finding is in agreement with our previous study in which we report an inverse relationship between BC and *L*_*i*_^37^.

### Effect of ligand binding on communication propensity

The methodology for calculating the communication/coordination propensity (CP) residue pair was first introduced for elastic network models by Chennubhotla and Bahar^55^ and subsequently extended by Morra et al.^33^ for MD trajectories. In context of the latter, CP for any two residues describes signal transduction events as a function of the mean-square distance fluctuation between the C_α_–C_α_ atoms over the trajectory. Inter-residue fluctuations that occur at lower intensities are expected to communicate more efficiently compared to that of high intensity C_α_–C_α_ fluctuations^33,55^. It is important to note that CP denotes communication time and thus smaller values represent more efficient commination between any two C_α_–C_α_ atoms. To obtain a compact representation of how bound ligands affect CP, we define the difference matrix between all residue pair CPs determined for the ligand-free and ligand bound systems (**Figure 8**). Thus, positive ΔCP (blue) indicates more efficient (faster) communication with the addition of ligand, while negative ΔCP (red) indicates slower communication (slower) in the presence of bound ligand.

**Figure 8:**
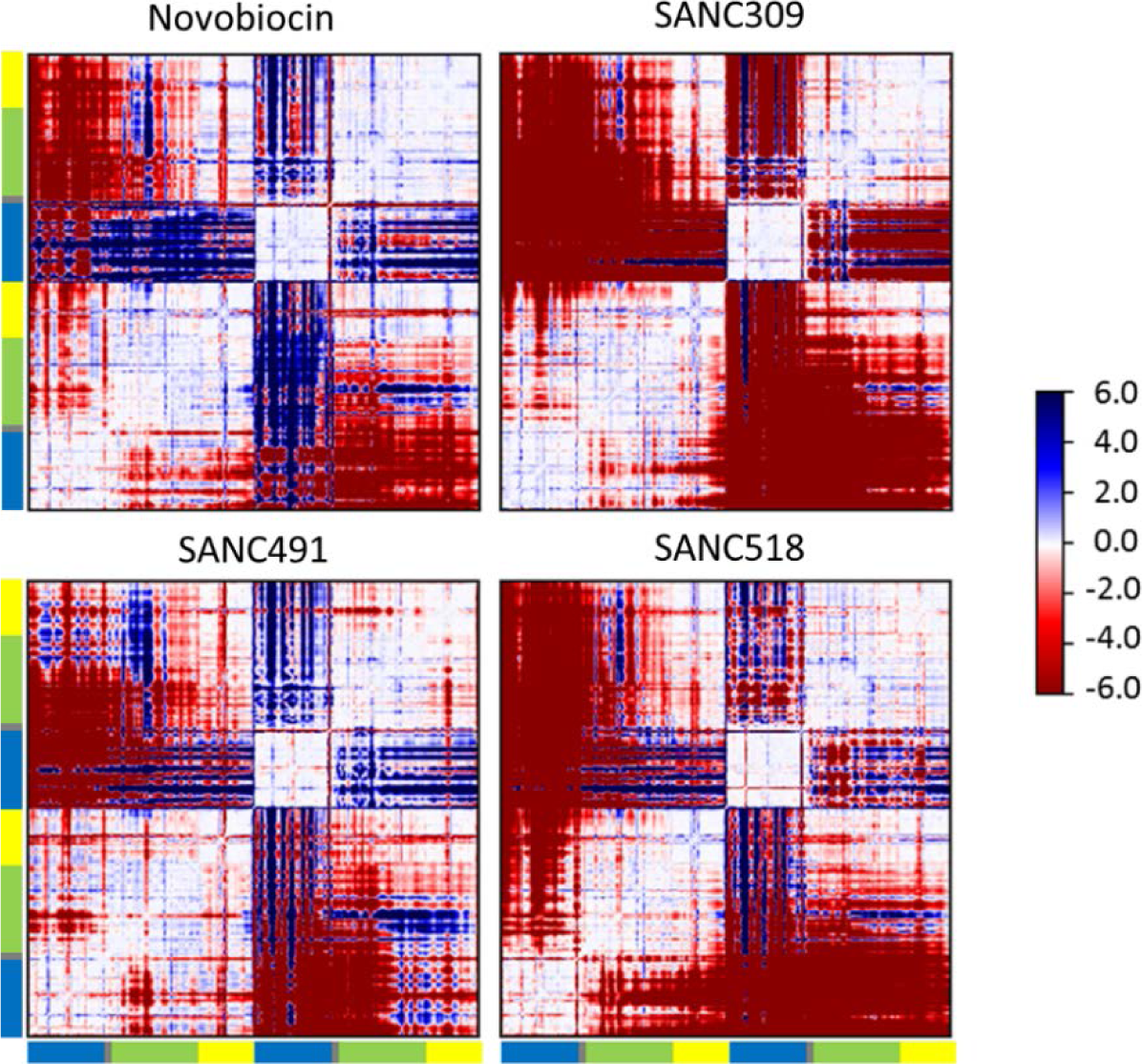
Difference in communication propensity. Matrices calculated as the difference in communication efficiency between the ligand-free complex and the ligand bound complex. Positive values (blue) denote higher communication propensity for the ligand bound complex compared to the ligand-free complex while negative values (red) indicate lower communication propensity for the ligand bound complexes.

Comparing the ΔCP matrices in **Figure 8**, we note a similar block character for regions of the protein that display either efficient (+CP) or slower (-CP) coordination propensities. The intra-protomer coordination patterns provide a visual means by which to compare the inter-domain communication propensities. For the Site-1 complexes, the NTD in protomer A appears to be decoupled from the M-domain and CTD as indicated by the red blocks. This decoupling is particularly apparent for SANC518 and resonates with our findings in the previous inter-domain contact analysis (**Figure 7**). In protomer B, the opposite effect is observed as indicated by the large blue blocks in which the NTD experiences more efficient CP between the M-domain and CTD, however this observation is less apparent for SANC518 which experiences some inter-domain decoupling in protomer B. Looking at the inter-protomer coordination patterns for these complexes, it is largely evident that ligand binding causes a loss in communication efficiency between protomers. However, in the presence of Novobiocin increased CP is observed between the NTD of protomer B and the M-domain:CTD region of protomer A, while SANC491 experiences a moderate increase in CP between at the NTD of protomer A and the CTD of protomer B. Looking at the coordination patterns SANC309, ligand binding at Site-2 appears to have a drastic effect on CP causing significant intra- and inter-domain decoupling within and between both protomers as evidenced by the large red blocks. It is interesting to note that this observation is particularly apparent in protomer B which provides the sole binding interactions for SANC309. Overall, these observations are consistent with the ΔRMSF results presented in **Figure 6**, whereby bound ligands at binding Site-1 appear to increase residue fluctuations at the NTD of protomer A and decrease fluctuations at the NTD of protomer B, while ligand binding at Site-2 increases residue fluctuations throughout both protomers.

We next investigate the relative contribution of individual residues to long range coordination with distant residues elsewhere in the protein. The average CP for neighbouring residues (*i* ± 4) for the ligand-free complex is 0.85, and we set CP=0.85 as a suitable threshold for discriminating fast communication between residues positioned >80 Å from one another in a similar manner described by Morra and co-workers^33^. By sequentially scanning the protein we record the fraction of residues in the whole protein capable of communicating with a CP ≤ 0.85 (**Figure 9**). Looking at the ligand-free complex it is evident that long range communication is established between the NTDs (residues 80-90, 150-160, 170-185) and CTDs (residues 555-580, 640-655, and 690-700) providing further evidence of NTD-CTD allosteric coupling in agreement with previous reports for yeast Hsp90 in the closed state^33^. For the ligand bound complexes, it is evident that none of the binding site residues directly contribute to long range residue communication (**Figure 9** – shaded areas). Rather, the presence and identity of the bound ligand appears to impact the relative long range coordination of residues in close proximity to the bound ligands. Novobiocin and SANC491 increase the fraction of communicating residues at NTD and CTD of both protomers (residues 80-90, 160-200, 555-580, 640-660 and 690-700). SANC309 appears to significantly reduce the communication efficiency of the CTD, as well as residues 80-90 at the NTD of protomer B, while SANC518 significantly reduces long range coordination in both protomers.

**Figure 9:**
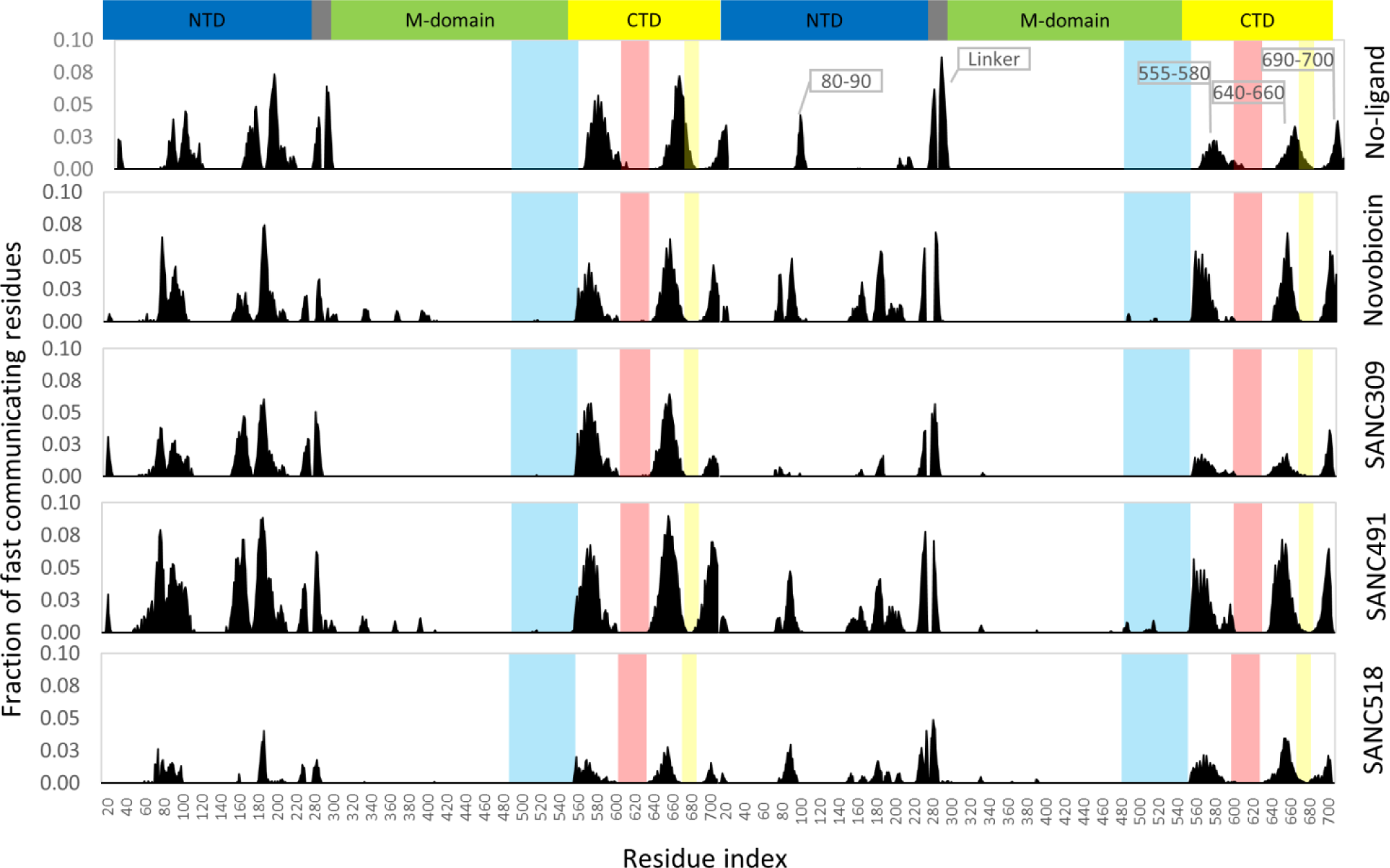
Fraction of fast communicating residues over 80 Å distances demonstrating communication efficiently. Each histogram refers to a single residue and indicates the total fraction of residues that communicate with it. Colored shading represents the ligand binding residues at Site-1 – yellow and red; and Site-2 – blue.

### Essential dynamics analysis

Given the tremendous size of Hsp90 (1400 residues), our 200 ns MD trajectories are of an insufficient length to access functional global motions such as the full closing motion expected for the ATP-only complex. To circumvent this, essential dynamics (ED) analysis techniques were employed to assess whether ligand binding corresponds to functional global correlated motions. In ED, MD trajectory is represented as the covariance matrix which is used to calculate eigenvectors and eigenvalues. The former metric represents the correlated displacement of atom groups through essential space, while the latter gives an indication of the magnitude (nm^2^). The configurational space represented by eigenvectors can be separated into two subspaces^56^; (i) the essential subspace which represents correlated motions comprising very few degrees of freedom, which likely point to functionally relevant global motions; and (ii) the independent subspace which is constrained to local regions and offer little functional importance. Here, we focus on the former subspace (i) which is often accounted for by the first few low-frequency modes with large corresponding eigenvalues. Analysis of the cumulative squared overlap for the first 20 modes for each complex (**Figure S5**), shows that between 60 % and 80 % of all protein motion is accounted for by the first three eigenvectors. In each case the first two eigenvectors account for more than 50 % of all protein fluctuations, suggesting these modes represent functionally relevant protein motions. Here, we illustrate the global correlated motions associated with the first eigenvector for each complex by projecting the corresponding trajectory onto the eigenvector and interpolating over the two most extreme projections and use arrows to describe the relative atomic displacements (**Figure 10**, **Figure S6** and **Movie S1-S5**).

**Figure 10:**
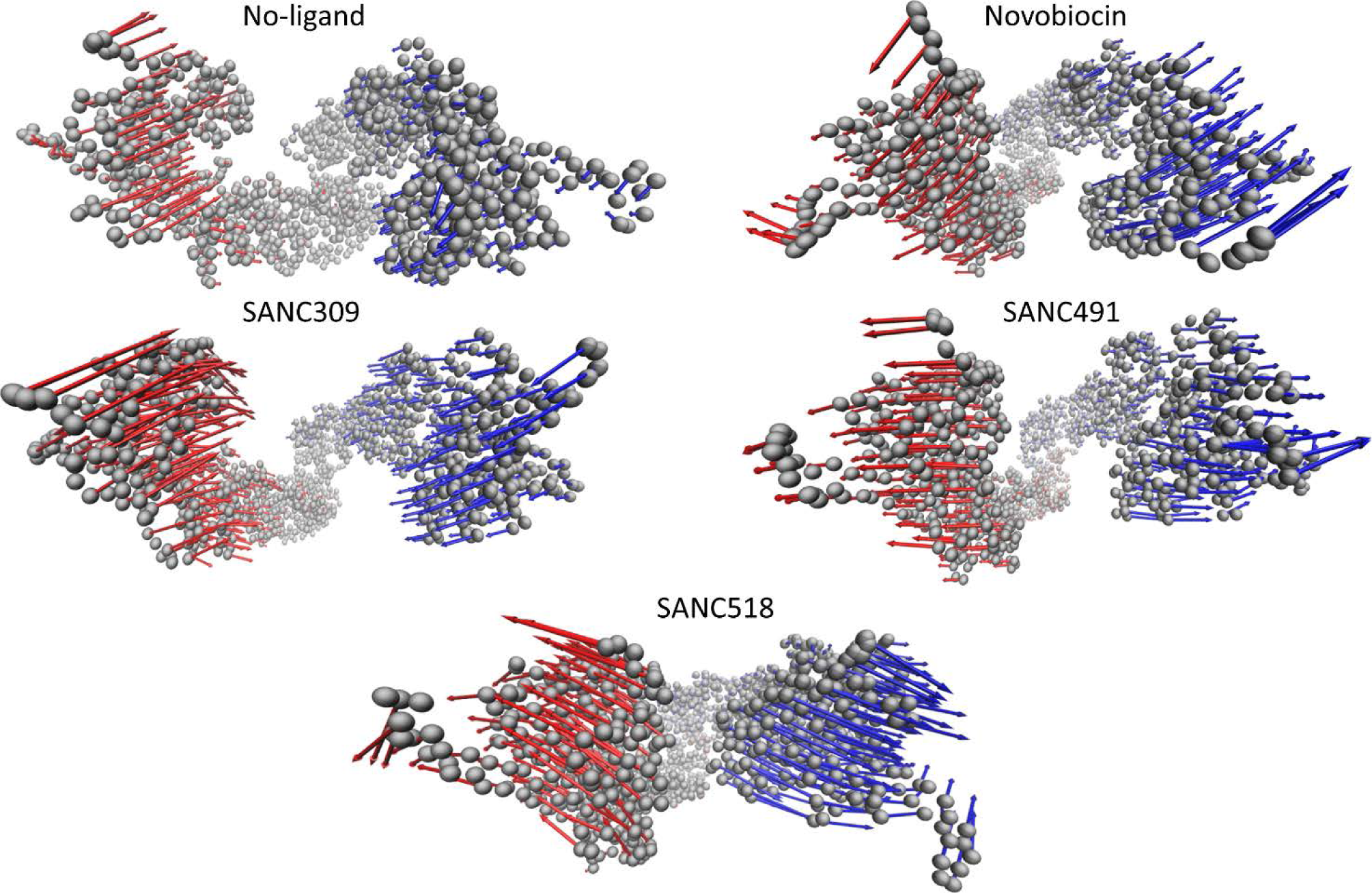
Top down view illustration of the essential dynamics displacements for the first eigenvector of each Hsp90 complex. The Hsp90 complex is represented by C_α_ atoms (grey spheres) and the arrows describe the relative direction and magnitude of the atomic displacements for protomer A (blue) and protomer B (red). Displacement arrows drawn for every 2^nd^ C_α_ atom to simplify presentation.

In the ligand-free state, ED analysis reveals linear atomic displacements for each protomer in the opposite direction towards one other, and closely resembles the closing transition, that is expected under these conditions, in which ATP binding triggers conformational rearrangements towards the catalytically active closed state. Addition of Novobiocin gives protomer displacements in opposite directions in an opening like motion. These displacements may provide early evidence of protomer uncoupling as Novobiocin is known to disrupt protomer dimerization at the CTD^32^. Interestingly, SANC491 and SANC518 demonstrate similar linear protomer displacements to Novobiocin, providing further evidence that these compounds may invoke a similar inhibitory allosteric mechanism. SANC309 on the other hand appears to enhance protomer displacements towards each other in a similar manner to the ATP-only complex. This observation coupled with larger displacements than the ligand-free complex, may suggest SANC309 to be an allosteric activator of human Hsp90α, by promoting allosteric closure of the dimer. Indeed, this observation agrees with earlier results in which SANC309 populated conformations with smaller inter-protomer distances (**Figure 4**-red). Furthermore, we note similar results for the second eigenvectors which describe rotational twisting of the protomers along their central axis in either a clockwise closing or an anti-clockwise opening motions much like the twisting and untwisting of a double stranded helix.

## Conclusions

Overall, our findings establish evidence of ligand specific modulation of the conformational dynamics of human Hsp90α in the open “v-like” conformation. We show how natural forces associated with ligand interactions at two putative druggable CTD binding sites direct the conformational sampling of the dimer by fine tuning the internal dynamics of the protein. Of the binding Site-1 ligands, Novobiocin and SANC491 enhanced conformational rigidity through protomer flexing and reduced NTD residue fluctuations resulting in more efficient NTD-CTD allosteric communication. SANC518, on the other hand, behaved differentially enhancing protomer flexibility leading to reduced NTD-CTD allosteric communication in protomer A. Given, the overlap in binding site of the Site-1 ligands with known inhibitors such as Novobiocin and several Bisphenol A based inhibitors, coupled with evidence of correlated opening motions, we propose that small molecules targeting the CTD dimerization site located at the four-helix bundle of the open conformation may externally modulate the conformational dynamics in favour of a more open conformation and thus act as allosteric inhibitors of Hsp90α. These compounds may prevent conformational cycling to the closed catalytically active state by either disrupting CTD dimerization as is the case for Novobiocin^32^, or by allosterically enhancing the energetic barrier that must be overcome in order to access the ATPase active state^23^. Finally, we note that Cephalostatin 17 (SANC491) is known to be a potent anti-cancer agent ^57,58^ although its mechanism of action remains unclear. In contrast, SANC309 at Site-2 greatly enhanced protein flexibility and decreased internal coordination, characteristics that may enable the protein to overcome the energetic requirements necessary to access the closed state and therefore act as an allosteric activator. Indeed, the correlated atomic displacements observed for this complex are in agreement with our previous PRS study, in which external force perturbations at Site-2 resulted in global protein displacements towards the closed catalytic conformation^37^, providing proof of concept for this approach through simulated ligand interactions. In summary, our findings provide novel insights regarding the selective external modulation of Hsp90α conformational dynamics, and shed valuable insight on allosteric drug development for Hsp90 as well as the protein’s complex allosteric mechanism of action. Additionally, this is the first study that applies combined information from PRS coupled with DRN to identify potential allosteric sites for inhibitor design. This proposed approach should be applicable in identifying allosteric drug targeting sites in other proteins.

## Methodology

### Molecular docking

The 3-dimensional coordinate data for human Hsp90α in an open “v-like” conformation was obtained from previous MD simulations of a homology model of human Hsp90 in the presence of various nucleotide conditions^37^. Clustering analysis was carried out on a 400 ns all-atom MD trajectory of Hsp90α in complex with ATP according to the methodology described by Daura and co-workers^59^, and the frame with the smallest RMSD to the average of the largest cluster selected the representative protein receptor. Natural compounds indigenous to Southern Africa were obtained from SANCDB^60^, while the known Hsp90 inhibitor Novobiocin was retrieved for the ZINC database. Both the compounds and the protein were prepared for docking using the AutoDockTools software suite^61^. The previously identified allosteric binding sites were mapped onto two separate grids with AutoGrid4^61^. Site-1 was centred over the four-helix bundle located at the CTD interface (residues G668-A685), and Site-2 on the adjacent pocket located on protomer B (residues H491-V508, V542-Q550, P596-N609). All docking calculations were performed with AutoDock4^61^ using the Lamarkian Genetic algorithm with a population size of 150, and the number of evaluations and generations set to 10 000 000. A total of 100 docking runs were performed for each compound, and each run evaluated with the semi-empirical scoring function supplied by AutoDock4. Clustering of the docked conformations was carried out using AutoDockTools with a cut-off of 1.5 Å. Docking of a compound was deemed reproducible if the largest cluster exceeded 50 % of the total runs, with an average energy score of less than −8.00 kcal/mol. The compounds were ranked by average energy score, and the lowest scoring conformation of the best candidates selected as the representative conformation for long range all-atom MD simulations.

### Molecular dynamics

All MD simulations were performed using GROMACS 5.1.2^62,63^ with in the CHARMM 36 force field^64–66^, using a orthorhombic periodic box with a clearance space of 1.5 nm. Water molecules were added as solvent, and modeled with the TIP3P water model, and the system neutralized using a 0.15 M NaCl concentration. Prior to production runs, each system was first energy minimized using a conjugate-gradient and energy relaxed up to 50 000 steps of steepest-descent energy minimization, and terminated when the maximum force < 1000.0 kJ/mol/nm. Energy minimization was followed by equilibration, first in the *NVT* ensemble at 310 K using Berendsen temperature coupling, and then in the *NPT* ensemble at 1 atm and 310 K until the desired average pressure (1 atm) was maintained and volumetric fluctuations stabilized. All production simulations were run for a minimum of 200 ns, and the backbone root-mean-square deviation (RMSD) of the protein monitored for convergence. Coordinate data for the protein and ligand were saved at 2 ps intervals for further analysis. All simulations utilized the LINCS algorithm for bond length constraints and the fast particle mesh Ewald method was used for long-range electrostatic forces. The switching function for van der Waals interactions was set to 1.0 nm and the cutoff to 1.2 nm. NH3^+^ and COO^−^ groups were used to achieve the zwitterionic form of the protein and periodic boundary conditions were applied in all directions and the simulation time step set to 2 fs. All trajectory clustering analyses were carried out with the GROMACS 5 cluster function using the gromos method described by Daura and co-workers^59^ using a cutoff of 0.2 nm. The RMSD values for each complex are displayed in **Figure S2**.

### Protein-ligand interactions

The Protein Ligand Interaction Profiler (PLIP)^17^ was used to predict protein-ligand interactions present over the course of each MD simulation. Using a sampling step of 0.1 ns, each trajectory was reduced to 2000 frames, and stored in PDB format. PLIP analysis was then performed on each frame, and in house Python scripts used to parse the PLIP interaction data for each time point and extract those residues involved in either hydrophobic or hydrogen bond interactions with the bound ligand. In this manner the time evolution of bond formation could be assessed.

### Dynamic residue networks

To analyse inter- and intra-domain communication, the protein is represented as a residue interaction network (RIN), where the C_β_ atoms of each residue (C_α_ for glycine) are treated as nodes within the network, and edges between nodes defined within a distance cut off of 6.7 Å^67^. In this manner, the RIN was constructed as a symmetric *N* × *N* matrix, where the *ij*^th^ element is assigned as 1 if residue *i* is connected to residue *j* and a zero if no connection exists.

In this study, MD-TASK^47^ was used to construct dynamic residue networks (DRN) for each MD trajectory, in which RINs are constructed for every *n*^th^ frame of the trajectory using a 200 ps time interval, to build a DRN matrix. By iterating over the DRN, each RIN is analysed in terms of the average of shortest path length (*L*_*ij*_) between residue *i* and any other residue *j*, and betweenness centrality (BC) of each residue. The shortest path length between two residues *i* and *j* is defined as being the number of nodes that need to be crossed to reach *j* from *i*. The average *L*_*ij*_ is then calculated as the average number of steps that the node/residue may be reached from all other residues in the RIN:

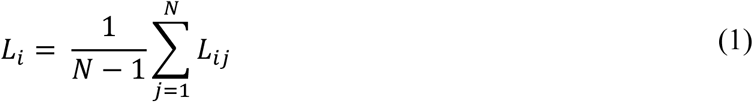

Here, we analyse the change in reachability (Δ*L*_*i*_) of each residue by monitoring how *L*_*i*_ shifts over the course of the MD trajectory:

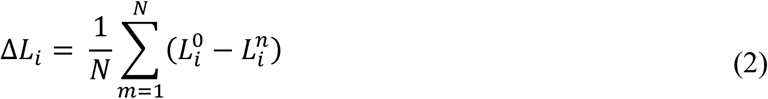

where 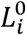 denotes the average shortest path length for residue *i* at time zero, and 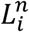 the average shortest path length at frame *n*

BC is defined as the number of shortest paths running through a node/residue for a given RIN, and provides a measure of usage frequency each node during navigation of the network. Here, BC was calculated using MD-TASK based on Dijkstra’s algorithm^68^ and the data rescaled in the range of 0.0 – 1.0 to be comparable between conformers. Finally, the average BC and Δ*L*_*i*_ are calculated over the DRN, as this measure provides an indication of residues that experience permanent changes in Δ*L*_*i*_ and BC as opposed to minor fluctuations over the course of the MD trajectory^47^ Inter-domain contacts between the NTD and M-domain were evaluated using contact_map.py script from MD-TASK with a distance cut-off of 6.7 Å. The script was utilized every 200 ps and the resultant data collated into a single data frame for plotting purposes.

### Communication propensity

The pairwise communication propensity (*CP*) describes the efficiency of communication between residues *i* and *j* and is based on the notion that signal transduction events in proteins are directly related to the distance fluctuation of the communicating atoms^33,55^. *CP* is thus defined as a function of the inter-residue distance fluctuations, where residues whose Cα -Cα distance fluctuates with low intensity are thought to communicate more efficiently (faster) compared to residues whose distance fluctuations are large in which the amplitude of the fluctuations results in slower inter-residue communication. *CP* is calculated as the mean-square fluctuation of the inter-residue distance

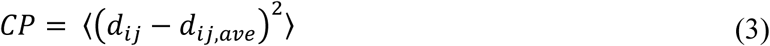

where *d*_*ij*_ is the time dependent distance between the Cα atoms of residues *i* and *j* respectively, and the brackets denote the time-average over the trajectory. In this study, the resultant *CP* matrix is used to assess the relative impact each compound has on the internal dynamics and overall flexibility of the protein, by calculating the difference matrix between each ligand bound complex to the ligand-free (ATP only) complex.

### Principal component analysis

Principal completed analysis (PCA) was used for the analysis of global motions present over the course of each MD simulation^56^. This technique involves two main steps: 1) the construction the covariance matrix, **C**, based on the positional deviation of each Cα atom, and 2) dimensionality reduction of **C** by diagonalization, to obtain the eigenvectors and eigenvalues. Each 3*N* x *N* covariance matrix was calculated based on an ensemble of protein structures obtained from the respective MD simulation and the elements of **C** defined as

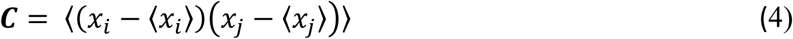

where *x*_*i*_ and *x*_*j*_ are atomic coordinates of each Cα atom, and the brackets denote the average. Eigenvectors with the largest eigenvalues are representative of the slowest modes, and generally are associated with large-scale movements in proteins, which are responsible for protein function. All PCA analyses were conducted using the GROMACS 5 software suite. The covar function was used for the construction and diagonalization of **C**, not including bound ligands and excluding the first 10 ns of trajectory data to avoid equilibration artefacts. The anaeig function was used to project the MD trajectories onto the main eigenvectors.

## Supporting information

**Figure S1: FTMap unbiased screen reveals two putative ligand binding sites in the CTD.** Site-1 is located at the four-helix bundle. Site-2 is mirrored in both protomers Site-2A and 2B respectively.

**Figure S2: RMSD plots for each ligand bound complex.** Black denotes the whole protein backbone RMSD, while blue and red denote the backbone RMSD of each individual protomer.

**Figure S3: Pearson’s correlation coefficients showing the relationship between RMSF, *L*_*i*_, and CP.** (**A**) *L*_*i*_ vs RMSF; (**B**) ΔRMSF vs Δ*L*_*i*_; (**C**) ΔCP vs ΔRMSF; (**D**) Δ*L*_*i*_ vs ΔCP.

**Figure S4: Change in betweenness centrality (ΔBC) for the ligand bound complexes relative to the ligand-free system.** Shaded regions denote binding site residues: blue sub-pocket, red helix_2_, yellow four-helix bundle. ★ Ligand interactions at the two ligand binding sites.

**Figure S5: (A) Cumulative squared overlap for the first 20 eigenvectors and (B) cosine content for the first eigenvector.**

**Figure S6: Side view illustrating the displacements of first eigenvector of each Hsp90 complex.** The Hsp90 complex is represented by C_α_ atoms (grey spheres) and the arrows describe the relative direction and magnitude of the atomic displacements for protomer A (blue) and protomer B (red). Displacement arrows drawn for every 2^nd^ C_α_ atom to simplify presentation.

**Movie S1: Correlated motions for the ligand-free complex:** Depicting dimer closure as protomer A (green) approaches protomer B (cyan).

**Movie S2: Correlated motions for the Novobiocin complex:** Depicting dimer opening as protomer A (green) and protomer B (cyan) displace in opposite directions.

**Movie S3: Correlated motions for the SANC309 complex:** Depicting dimer closure as protomer A (green) approaches protomer B (cyan).

**Movie S4: Correlated motions for the SANC491complex:** Depicting dimer opening as protomer A (green) and protomer B (cyan) displace in opposite direction.

**Movie S5: Correlated motions for the SANC518 complex:** Depicting dimer opening as protomer A (green) and protomer B (cyan) displace in opposite directions.

## Acknowledgements

This work is supported by the National Research Foundation (NRF) South Africa (Grant Number 93690). The content of this publication is solely the responsibility of the authors and does not necessarily represent the official views of the funder. The authors thank the Centre for High Performance Computing (CHPC), South Africa, for computing resources.

## Author Contributions

DP and ÖTB conceived the study. DP designed the study details, wrote the scripts, and did the calculations and analysis under the supervision of ÖTB. DP wrote the initial draft manuscript. ÖTB and DP revised and contributed towards the final manuscript.

